# A Novel Rapidly Manufacturable Flexible Subdural Electrode Array for Intraoperative Mapping of Cortical Activity

**DOI:** 10.64898/2026.03.05.709791

**Authors:** A.R. Mamleev, D.S. Suchkov, E.I. Malyshev, A.A. Vorobyov, V.R. Sitdikova, V.M. Silaeva, A.E. Logashkin, A.K. Kireev, M.A. Sorokina, D.M. Mitin, I.S. Mukhin, V.V. Belousov

## Abstract

Flexible and biocompatible neurointerfaces are crucial elements for intraoperative monitoring and chronic neural recordings. However, existing fabrication methods often involve complex cleanroom processes, limiting rapid prototyping and customization. In this study, we present a fast, low-cost method for manufacturing a flexible subdural electrode array based on polydimethylsiloxane (PDMS) and gold conductive layer. The fabrication process utilizes a laser cutter for both mask generation and direct patterning of metal traces on a PDMS substrate, achieving a resolution of up to 30 *µ*m. A detachable interface was developed for reliable connectivity during testing. The electrochemical and mechanical properties of the array were characterized, demonstrating Ohmic behavior and stable conductivity after 50 cycles of mechanical bending, with a degradation of less than 10%. Electrochemical impedance spectroscopy (EIS) confirmed the viability of the electrodes for recording physiological signals. The functionality of the array was validated *in vivo* by performing simultaneous recordings of local field potentials (LFPs) and electrocorticography (ECoG) in the rat somatosensory cortex. The signals from the flexible subdural array showed a statistically significant (*p* < 0.001) median cross-correlation of 0.35 with LFPs recorded at a depth of 600–800 *µ*m by industrial electrode. We demonstrate here a robust and accessible approach for producing functional neural interfaces, suitable for rapid iteration and customization in research and clinical applications.

## Introduction

The functionality of our brain is the result of the activity of neural networks consisting of thousands of neurons. Neuronal activity is determined by the generation of action potentials (APs) across the neuronal membrane, each of which is triggered by the spatiotemporal summation of synaptic input signals onto the neuron. [41]. To study this activity, researchers have developed various types of electrophysiological recording methods, divided into extracellular and intracellular methods. Extracellular recording allows one to obtain data on the overall activity of the neural ensemble surrounding the recording electrodes. An array of extracellular recording electrodes of varying configurations forms an electrode matrix, which serves as the communication interface between neurons and the recording system. Data obtained from such electrode matrices can be used to analyze the recorded neural signals for visualizing brain activity or implementing control algorithms for robotic limbs. However, this is only achievable when recording signals with a high signal-to-noise ratio (SNR) and relative biocompatibility of the electrode matrix materials with brain tissue. The parameters of electrode matrices are a fundamental factor determining their effectiveness in key areas of applied medicine.

Electrode arrays *in vivo* have different geometric and electrophysical parameters, as well as manufacturing materials, depending on their location relative to the brain during signal recording. When selecting a suitable electrode array, it is necessary to consider the trade-off between the quality of the recorded signal and the degree of invasiveness. For example, electrode arrays implanted in the brain have higher signal-to-noise ratios (SNR) than EEG arrays placed on the scalp. Currently, electrode arrays based on polymers with a low Young’s modulus are preferred. The material used to manufacture electrode arrays is a key aspect of their functionality, so one of the main goals of modern research is to increase their flexibility and elasticity. Research in the field of flexible electronics began approximately 30 years ago [3, 13] in response to the demand for macroelectronics [42], such as flexible light-emitting devices [15, 44]. Flexible and stretchable electronics demonstrated their potential applications in the late 2000s when the concept of biologically integrated electronics was proposed [45]. This paved the way for the creation of stable and long-lasting personalized bioelectronic interfaces, including epidermal devices for vital sign monitoring, [15, 24, 45], brain-computer interfaces with devices for electrocardiogram, electrocorticography, and electromyography mapping [23, 26, 52, 53, 55], as well as minimally invasive surgical instruments [25, 27]. When using such bioelectronic interfaces, it is crucial that they match the mechanical properties of the biological system’s tissues, ensuring biocompatibility. For brain-computer interfaces, electrode arrays must, firstly, detect changes in electrical potential in the range of several tens of microvolts; secondly, they must have high spatial resolution; thirdly, the substrate on which the electrode array is placed must be thin enough to ensure the absence of inelastic mechanical interaction with the brain; and finally, the electrode array material must be biocompatible. It has been shown that such bioelectronic interfaces can be further integrated with wireless signal transmission systems such as Bluetooth and NFC technologies [33, 34], and the ability to function in an aquatic environment has also been demonstrated [32]. Due to their high portability and adaptability, bioelectronic interfaces are also successfully used in systems for diagnosing the state of the body [37].

Advances in materials science have led to the development of high-density planar non-penetrating electrode arrays for electroencephalography and electrocorticography. However, measuring biological electrical signals using electrode arrays must not cause harm to the human body. Therefore, it is recommended that the substrate and metal components comprising the electrode be made of materials that do not cause chronic reactions, such as those caused by the immune system. When developing flexible electrode arrays for implantation, special attention is paid to long-term biocompatibility and biostability under real-world recording conditions.

The structure of any electrode array is conventionally divided into a substrate and an electrode portion. If the electrode array substrate is too rigid, it can damage brain tissue, whose Young’s modulus is approximately 3 kPa. Therefore, it is crucial to use a material with an appropriate Young’s modulus, close to that of brain tissue. For silicon used in microelectrodes, this value is close to 130–185 GPa. The Young’s modulus of polymeric materials is typically in the range of several GPa—significantly lower than that of silicon, but still higher than that of brain tissue. Such materials include, for example, the biocompatible photopolymer SU-8 [2, 7], polydimethylsiloxane (PDMS) [29], polyimide (PI) [31], and poly(chloro-p-xylylene) (Parylene C) [18, 48]. The brain-like Young’s modulus of these materials makes them suitable for use as substrates for electrode arrays. Shape-memory polymers, such as (parylene-C + polyvinyl alcohol) (PVA) [49], liquid crystal polymers [19], and composites based on cellulose nanofibers and a polyvinyl acetate matrix [17], are considered to be a promising materials. They could reduce invasiveness by better matching the Young’s modulus of brain tissue and the electrode array substrate material. Currently used electrode arrays allow for stable measurement of brain electrical signals *in vivo* over several days or weeks [1, 5, 20, 40, 41]. Electrode arrays fabricated using SU-8 as a substrate were also shown to successfully record action potentials of individual neurons for 8 months. [12].

The ultimate goal of creating electrode arrays is to safely record electrical signals from the brain over long periods of time. The material used in the conductive portion of the electrode arrays must have high electrical conductivity for signal transmission while being non-toxic to living organisms. Copper (Cu) [6], gold (Au) [28, 30, 47], platinum (Pt) [8, 21, 36, 39, 51, 54], silver (Ag) [10], titanium (Ti) [4], tungsten (W) [43], indium tin oxide [56] and graphene [38] can be used as promising conductive materials for forming electrodes and contact pads, as they possess high electrical conductivity at room temperature. However, copper and silver have toxic effects on brain tissue when exposed to long-term exposure, making them unsuitable for use in chronic implantation. At the same time, platinum, gold, or materials such as highly doped polysilicon are considered safe for long-term exposure [14, 46]. Recently, conductive polymers have been used as non-toxic materials, an example of which is poly(3,4-ethylenedioxythiophene) (PEDOT) [22, 35], polypyrrole (PPy) [16], and polyaniline (PANI) [9]. Thus, the use of biocompatible polymer-based neural devices is a promising direction of development.

## Materials and Methods

### 0.1 Ethical Approval

The study was conducted in accordance with Ethical commitee of the Pirogov Russian National Research Medical University (ethical approvals 02/2025 and 02A/2025). The described surgical procedures and animal handling methods were previously validated by the local ethics committees of several Russian and international organizations. Experiments were conducted in accordance with the “Guide for the Care and Use of Laboratory Animals,” National Academy Press (2011), as well as with Directive 2010/63/EU of the European Parliament and of the Council of the European Union of 22 September 2010 on the protection of animals used for scientific purposes (Article 27).

### Electrode Fabrication

We developed a prototype flexible subdural matrix for intraoperative monitoring. This matrix is based on a polydimethylsiloxane (PDMS) substrate with gold conductors (Figure1A, left; Figure1C). The fabrication technology consists of four main steps: (1) manufacturing a photomask, (2) creating a blank on a glass carrier substrate with a first PDMS layer and an applied conductive layer, (3) etching conductive tracks, and (4) forming a dielectric layer and exposing contact pads and electrodes.

**Figure 1.**
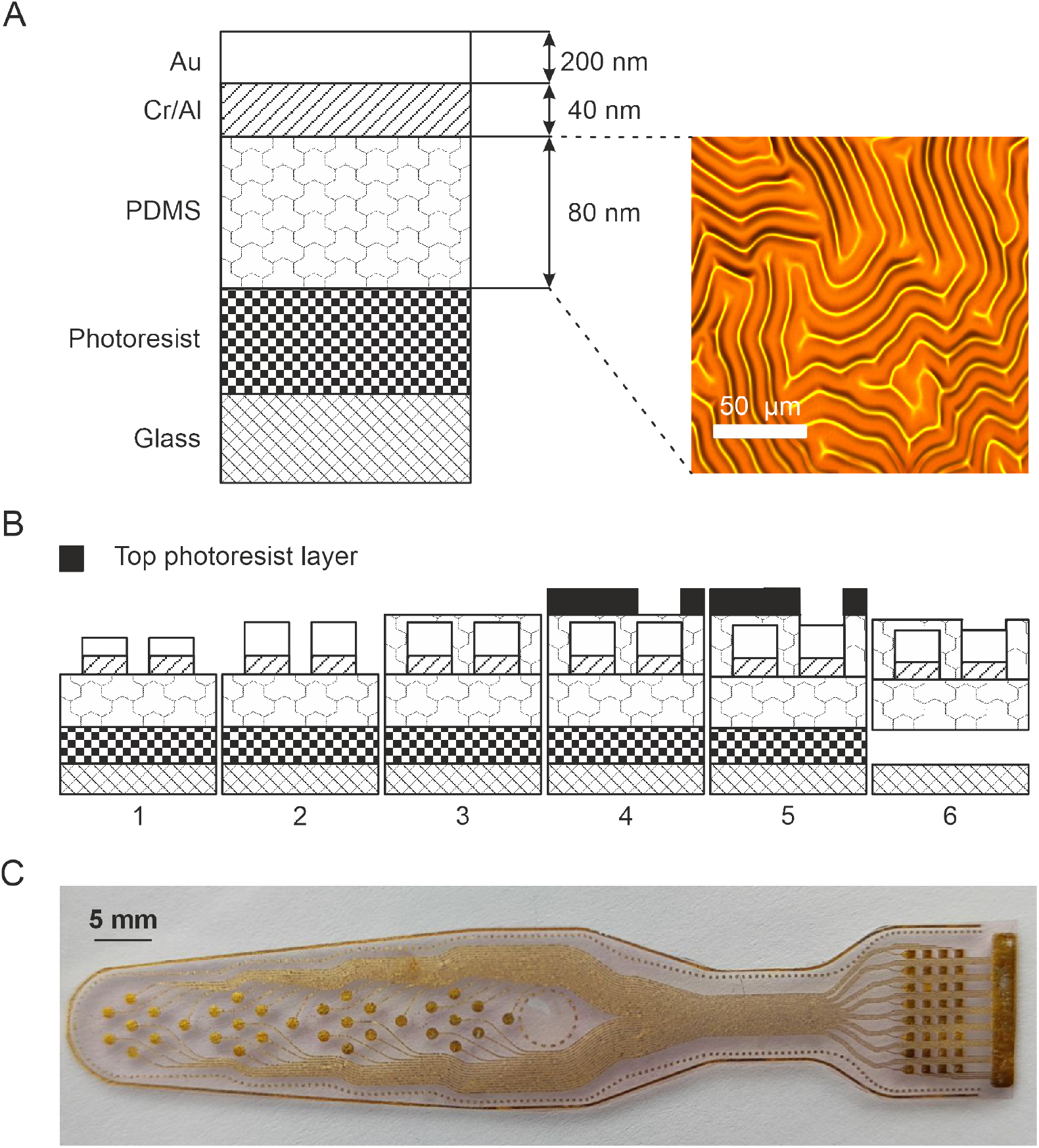
Schematic illustration of the fabrication process for the flexible subdural electrode array. **A**, Structure of the initial electrode array template (left), consisting of sequentially deposited layers (from bottom to top): glass, photoresist, polydimethylsiloxane (PDMS), Cr/Al layer, Au layer. Scanning electron microscope image of the multielectrode array elements on a flexible PDMS substrate (right). **B**, Schematic of the main stages of forming the multielectrode array on a flexible substrate: (1) Formation of the conductive pattern (Au + Cr/Al) by laser engraving; (2) Galvanization of conductive elements; (3) Encapsulation of conductive elements with a monolithic PDMS layer (black); (4) Application of a top photoresist layer and its exposure through a photomask; (5) Plasma-chemical etching of PDMS in areas unprotected by photoresist; Removal of the top and sacrificial photoresist layers. **C**, Photograph of the fabricated flexible subdural electrode array.

The first step is to fabricate a photomask for subsequent lithography. Direct laser engraving was used for this purpose, ensuring the required accuracy (approximately 10 µm) at high throughput. Optical glass, thoroughly cleaned in a soda solution and deionized water, was used as a substrate, then activated in low-temperature oxygen plasma. A continuous layer of aluminum was deposited on the prepared surface in a vacuum system. The pattern was formed using a laser erosion system in ultrashort pulse mode, ensuring clean metal removal without damaging the substrate or creating burrs. After engraving, the photomask was cleaned with compressed air and rinsed with deionized water.

The next step is to create a preform, where a glass substrate serves as a temporary rigid support. It is sequentially cleaned in a soda solution, deionized water, and oxygen plasma to achieve high levels of purity and adhesion. A layer of soluble photoresist is then deposited on the glass, serving as a sacrificial layer for subsequent separation of the finished device. Liquid PDMS is then spin-coated, which, after polymerization at 60^*°*^C for 12 hours, forms the flexible substrate of the array. This temperature selection ensures the formation of a microrelief on the PDMS surface (Figure1A, right) upon cooling. This, combined with thermal deformations in subsequent processes, significantly improves the mechanical stability of the final product. A multilayer conductive layer consisting of an adhesive chromium sublayer, an aluminum layer, and a base conductive gold layer is deposited on the cured PDMS in a vacuum system using thermal evaporation. (Figure1A, left).

The formation of electrodes and conductive paths is carried out by direct laser ablation. (Figure1B). The engraving process is performed on a laser erosion system in a gentle mode using ultrashort pulses lasting approximately 8 nanoseconds. This enables selective and precise metal evaporation, forming tracks with a resolution of up to 10 micrometers while minimizing the thermal load on the sensitive PDMS substrate, preventing damage and melting of the conductor edges. Further, to increase the mechanical strength of the conductive pattern, it was created using galvanic gold deposition. (Figure1B).

The final step involves forming a top insulating layer and providing electrical access to the contact areas. To completely isolate the conductive paths, the entire surface of the structure is covered with a second layer of PDMS (Figure1B), which, after polymerization, creates a biocompatible and flexible dielectric barrier. To expose the contact pads and electrodes, photoresist is applied to this layer. The photoresist is exposed through a prefabricated photomask and developed, forming a protective mask (Figure1B). Then, plasma-chemical etching in an oxygen-sulfur fluoride atmosphere selectively removes the PDMS in the unprotected windows of the mask, revealing the underlying metal electrode (Figure1B). After removing the mask, the completed flexible matrix is separated from the glass substrate by dissolving the sacrificial photoresist layer in a special solvent (Figure1B).

A prototype of a removable interface for the array was also developed (Figure3A). This interface provides a reliable, removable electrical connection and mechanical fixation between the implantable portion of the array and peripheral equipment, such as a stimulator or recorder. It is intended for quality control of implantable parts during manufacturing (specifically, for impedance measurements) and for conducting laboratory experiments on animals.

### Interface

Structurally, the detachable interface consists of two key components:

1. **Interface Board:** Fabricated on a printed circuit board (PCB), this component features an industrial connector (MOLC-110-01-S-Q, manufactured by Samtec) on one side (Figure3A) for connection to peripheral equipment. On the opposite side are spring-loaded contacts (pogo pins). These contacts provide a reliable, multi-cycle, releasable electrical connection to the implanted portion of the subdural electrode, ensuring that the contact pads on the implanted portion remain intact.
2. **Clamping Mechanism:** This interface connector element serves two main functions. First, it ensures reliable mechanical contact between the spring-loaded contacts of the interface board and the contact pads of the implanted section. Second, it firmly secures the implanted section of the subdural electrode relative to the interface board at the connection point, thereby preventing any displacement.

### Electrical and Mechanical Characterization

Tests were conducted to investigate the properties of the developed prototype. To study conductivity, the current-voltage characteristics (CVC) of individual conductive traces and contact pads were measured. The measurements were performed using a precision source-measurement instrument (Keithley 2400) operating in constant-current mode while simultaneously monitoring the voltage drop. The resulting linear dependences confirmed the ohmic nature of the contacts (Figure2A). Surface resistance was calculated using the standard two-probe method. The resistance of the current path material was less than 100 Ω/sq.

**Figure 2.**
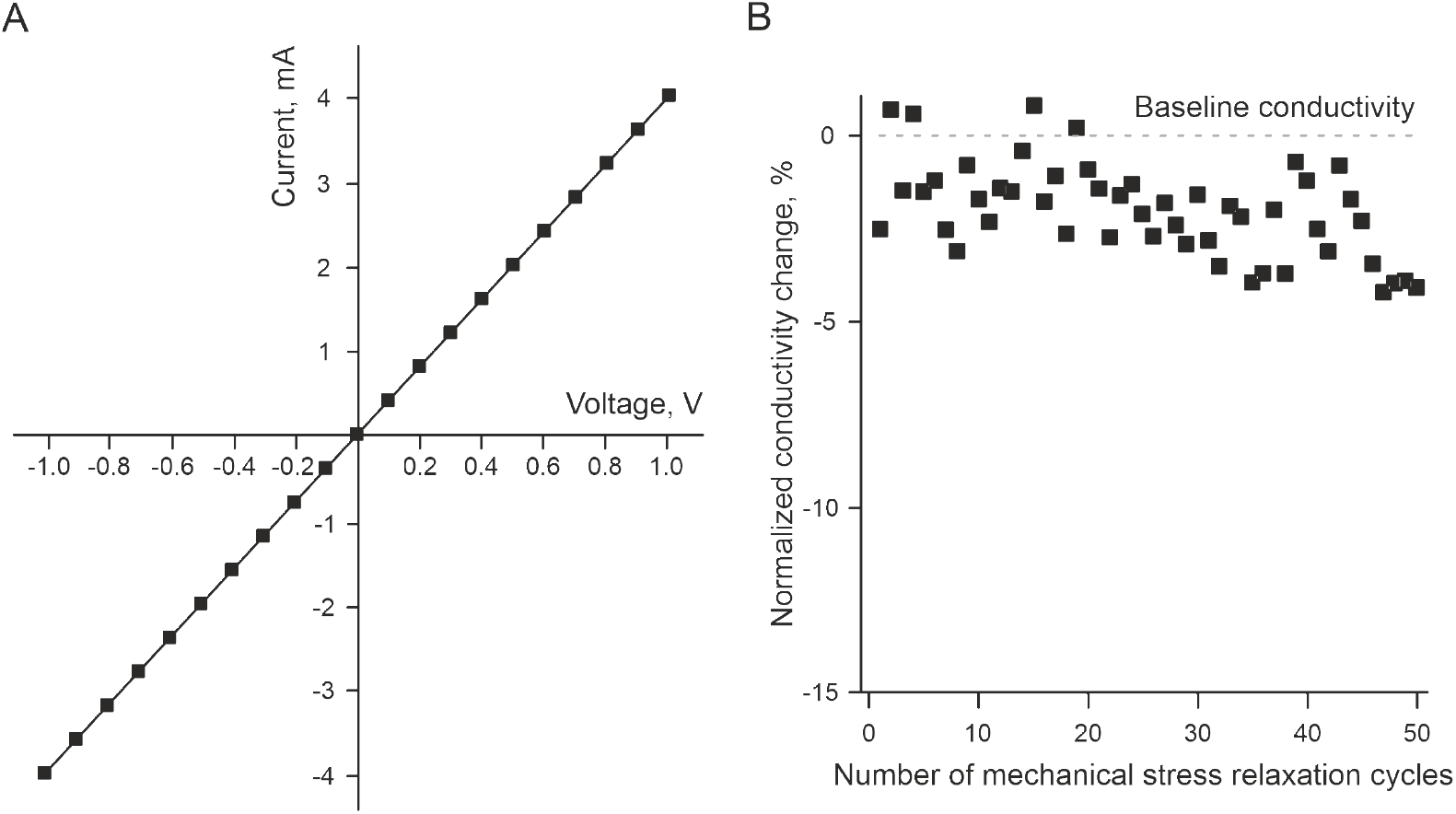
Electrical properties of the flexible subdural electrode. **A**, Currentvoltage (I-V) characteristic of a conductive trace fabricated on a flexible substrate. **B**, Results of measuring the relative change in conductivity of a conductive trace during cycles of bending and relaxation of applied mechanical stress.

To assess mechanical reliability and durability, a specialized study was conducted to assess the stability of electrophysiological parameters under cyclic deformation. The experiment included a series of impedance and conductance measurements before and after a specified number of mechanical deformation cycles around a cylindrical mandrel with a controlled radius of curvature. The measurement procedure was repeated after each stage of mechanical loading to determine degradation of key characteristics. (Figure2B).

To comprehensively evaluate the frequency characteristics and state of the electrode-electrolyte interface, impedance spectroscopy was performed using a NanoZ impedance analyzer (Plexon Inc, USA). Measurements were performed in a standard two-electrode configuration, where the electrode under study was characterized relative to a silver/silver chloride (Ag/AgCl) reference electrode in a phosphate-buffered saline (PBS, pH 7.4). These measurements yielded data on the dependence of the flexible subdural electrode impedance on the applied signal frequency (from 1 Hz to 5 kHz) (Figure3C).

**Figure 3.**
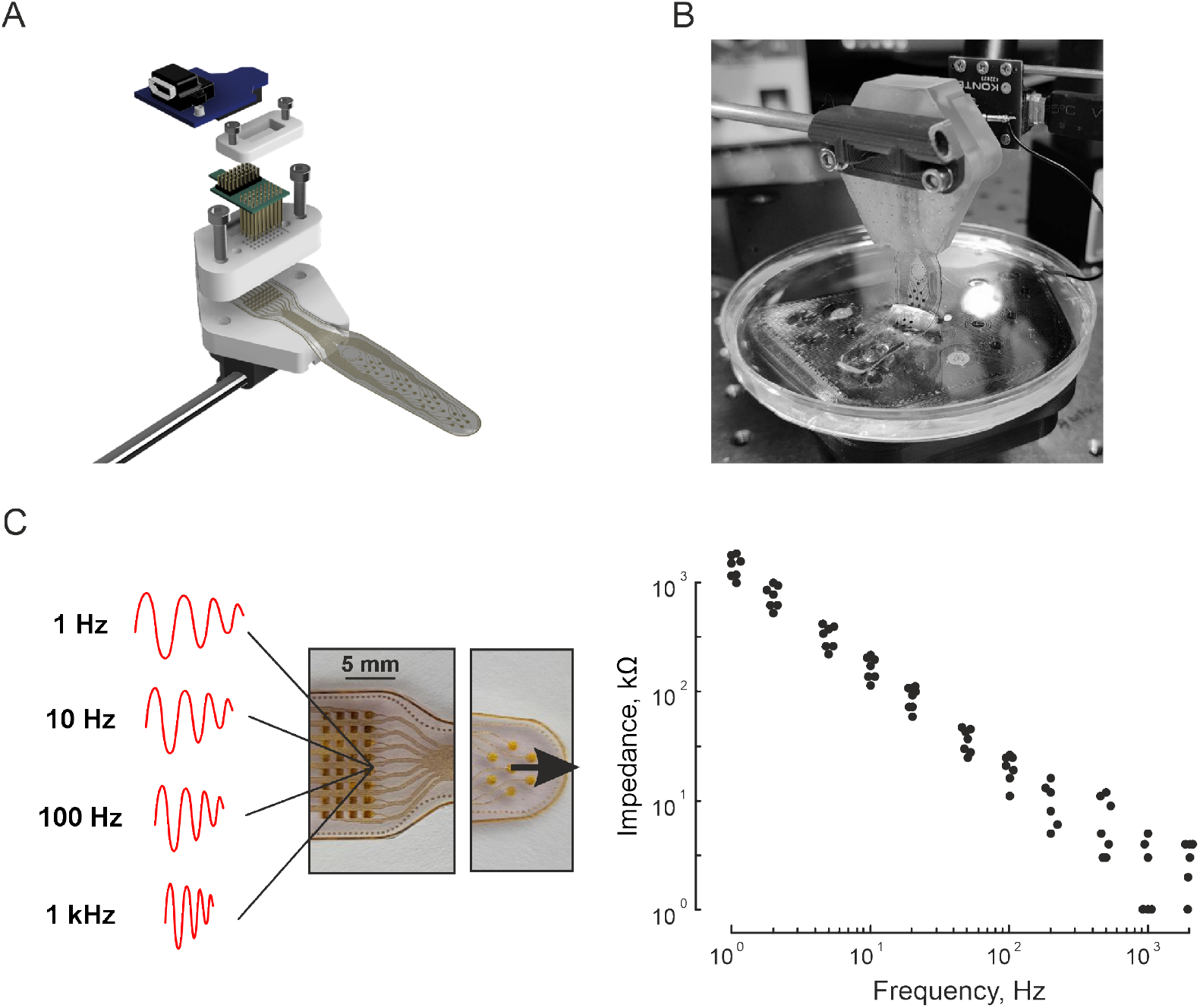
Impedance spectroscopy of the flexible subdural electrode. **A**, Schematic of the developed interface for connection to the subdural array. **B**, Photograph of the impedance measurement setup. **C**, Impedance spectroscopy results.

### *In Vivo* Validation: Surgical Procedure and Electrophysiological Recordings

To test the prototype’s functionality, animal experiments were conducted recording brain electrical activity. The experiments were performed on Wistar rats of both sexes aged 60 to 74 days. Surgery was performed under anesthesia using gaseous isoflurane (Baxter, USA) (5% for induction, 1.5% for maintenance). For additional local analgesia, a 2% lidocaine solution (1 mg/kg) was administered subcutaneously. During surgery, the skin was removed from the dorsal surface of the skull. Dental cement was applied to the cleaned and dried bone surface to secure the superfusion chamber, the bottom of which was the skull surface. For subsequent anesthesia, urethane (1.5 g/kg, Sigma-Aldrich, USA) was administered intraperitoneally.

To stabilize and immobilize the animal’s head, the edges of the superfusion chamber were fixed on a stereotaxic apparatus (Figure4A). The bone in the pre-marked area was then sequentially thinned with a high-speed drill to a translucent state under constant perfusion with isotonic saline (154 mM NaCl). The perimeter of the thinned area was carefully separated from the dura mater (DMA) using surgical tweezers. The DMA region over the target cortical area was removed. During the experiment, the animal was additionally warmed to a physiological temperature of 35–37^*°*^C using a heating pad.

**Figure 4.**
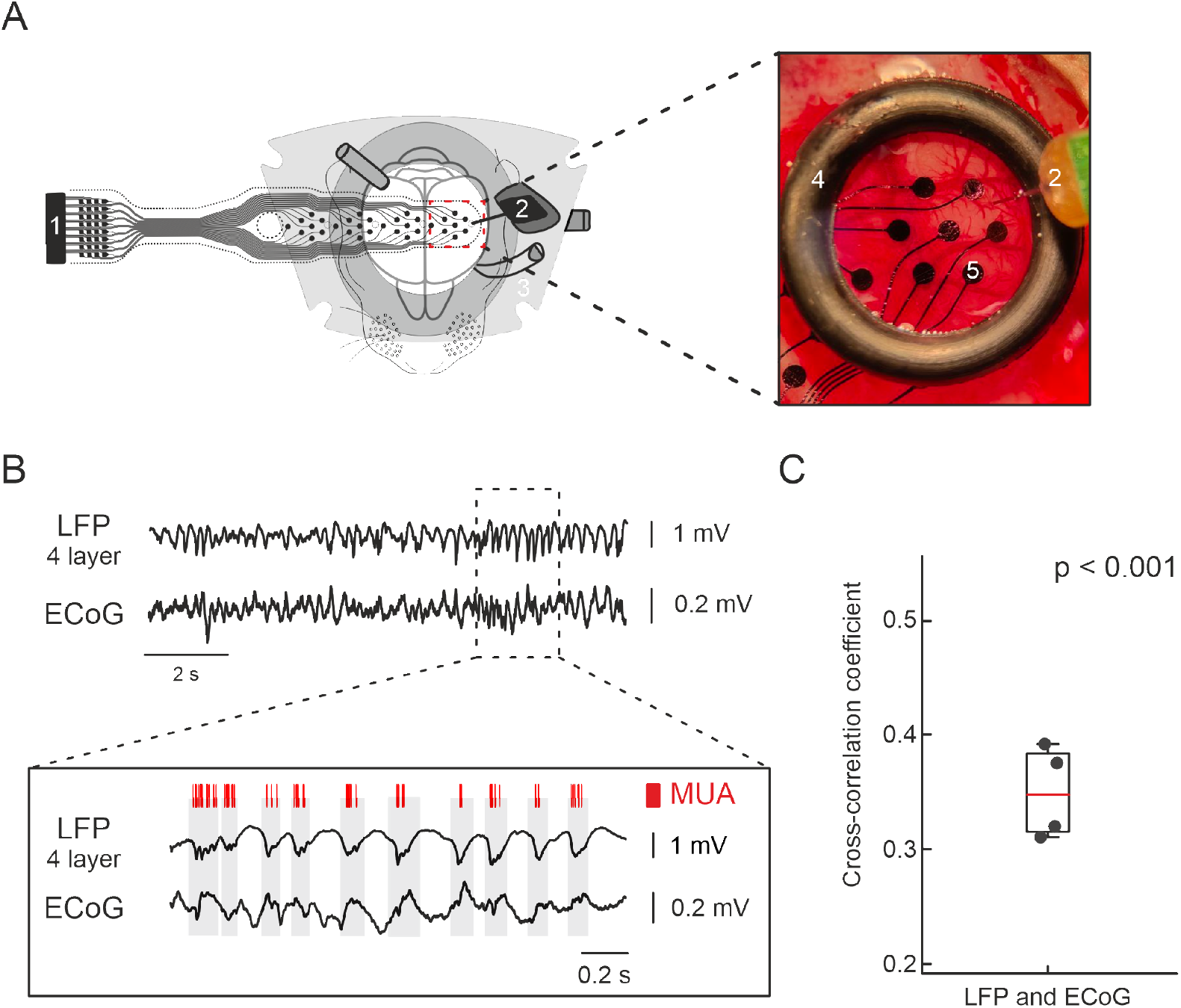
Data of the flexible subdural array for intraoperative monitoring. **A**, Schematic of the acute experiment setup for simultaneous recording of rat brain electrical activity using the flexible subdural array and a penetrating electrode (left), and a micrograph of the in vivo electrode and array placement (right): 1 – subdural electrode array; 2 – invasive depth electrode; 3 – superfusion chamber; 4 – black retaining ring; 5 – electrode sites of the subdural array. **B**, Example of simultaneous extracellular recordings: local field potential from 4 layer of somatosensory cotrex(LFP, top trace), subdural ECoG (middle trace), and an expanded window showing an example of slow-wave synchronous oscillations (bottom trace). Red ticks indicate multiunit activity (MUA) during periods of neuronal activity (gray boxes). **C** Group statistics results for the cross-correlation between the two signals (LFP-layer 4 and ECoG, *p − value <* 0.001).

Cortical activity was recorded in the somatosensory system using a 16-channel silicon-based electrode (100 *µ*m pitch between recording sites, Neuronexus Technologies, USA) (Figure4A). The vertical arrangement of electrodes allowed simultaneous recording of cortical activity from different layers of the same cortical column.

Electrocorticography (ECoG) was performed using the developed flexible subdural array prototype. This array was placed on the surface such that the electrode sites were located on each hemisphere and in direct contact with the brain surface (Figure4A). A silver/silver chloride electrode placed in the solution bathing the neocortex surface served as the reference. During the experiment, recordings of local field potentials (LFPs) and ECoG were performed (Figure4B). The signal recorded by the electrode was amplified and filtered (*×*192; 0.1 Hz to 7 kHz) using an XDAQ CORE2 amplifier (KonteX neuroscience, Taiwan), and recorded at a sampling rate of 30,000 Hz.

### Data Analysis

Data analysis was performed using the wEEGit software package [50] and custom functions written in the Matlab environment. At the first stage, multiunit activity (MUA) was detected. For this, the signal was filtered (300–6000 Hz); all negative events below a threshold of 10 standard deviations were considered MUA (Figure4B). At the second stage, the cross-correlation was calculated using the normalized cross-correlation function between the signal from the dipole electrode channel and the signal from the subdural array electrode located in the same area of the cerebral cortex. The median cross-correlation coefficient between potential fluctuations in the subdural array area and the local field potential at a depth of 600–800 *µ*m in the somatosensory cortex was 0.35 (25% – 0.32 and 75% – 0.38, n = 4 animals). Data are presented as median with interquartile range (Figure4C). To assess the statistical significance of the cross-correlation between signals, a non-parametric permutation test was applied. Permutation was performed by randomly shuffling the timestamps of one of the signals using a random permutation algorithm (MatLab function randperm). For each case, the maximum absolute cross-correlation within the range of allowable time shifts was calculated. The significance level was determined as the proportion of permutations (n = 1000) in which the maximum absolute correlation exceeded the observed value. The threshold significance level was set at *α* = 0.001.

## Results

### Fabrication and Basic Characteristics

The process described above has proven to be highly reliable for the fabrication of flexible subdural arrays with laser ablation-defined electrode geometry. As shown in Figure1A (right), the process resulted in well-defined structures. Conductive tracks, composed of a multilayer Au/Cr/Al structure, demonstrated excellent adhesion to the PDMS substrate throughout all subsequent processing steps, with no signs of delamination or cracking. This strong adhesion is attributed to the presence of a chromium adhesion layer and optimized, gentle laser engraving parameters, which minimized thermal stress at the metal-polymer interface.

### Electrical and Mechanical Properties

The current-voltage characteristics of the conductive traces were linear (Figure2A), confirming Ohmic behavior and the integrity of the metallic conductors. The Resistivity of the material of the current traces, fabricated using photolithography on solid substrates, was no less than 100 Ω/sq. The measured I-V curve (Figure2A) demostrated a linear dependence of the conductive paths.

Mechanical cycling tests revealed the robustness of the array. After 50 cycles of bending and relaxation of mechanical stresses, the change in conductivity was less than 10% of the initial value (relative to the initial conductivity before bending, *I*_initial_*/I*_bent_). Conductivity typically decreased upon bending and returned to near-initial values upon relaxation (Figure2B), indicating good mechanical resilience without catastrophic failure of the traces.Electrochemical impedance spectroscopy (EIS) provided the frequency-dependent impedance profile of the electrodes (Figure3C). The data showed the characteristic decrease in impedance with increasing frequency, typical for an electrode-electrolyte interface, confirming their suitability for recording bioelectrical signals in the relevant frequency bands for ECoG and LFP.

### *In Vivo* Validation

The flexible array was successfully implanted in the subdural space over the somatosensory cortex (Figure4A). During experimental studies on rats, despite a total experiment duration of up to 6 hours, no cases of perioperative mortality or unstable physiological state were recorded. Throughout the entire observation period, we were able to obtain stable and reproducible recordings of spontaneous cortical activity using the tested arrays—the lengthy nature of the experiment did not lead to deterioration in signal quality or the appearance of significant artifacts associated with fatigue during preparation. Careful movement or replacement of arrays on the surface of the cerebral cortex did not cause bleeding or alter the quality of the recorded signal, provided the cortex was constantly hydrated with saline. Successful surgical access to the rat cerebral cortex required precise cranial trepanation, taking into account the delicate bone structure and the need to preserve the integrity of the dura mater before implantation. Key factors included meticulous hemostasis, prevention of cerebral edema, and fixation of the animal’s head in a stereotaxic apparatus to minimize respiratory and pulse artifacts during long-term recording.

Simultaneous recording with a penetrating silicon probe allowed direct comparison of signals recorded by the surface array (ECoG) and the local field potential (LFP) from the cortical layers (Figure4B). Clear neurophysiological signals, including slow oscillations, were observed in both recording modes.

Quantitative analysis revealed a significant correlation between the cortical activity registered by the two interfaces. The median cross-correlation coefficient between the ECoG signal from the flexible array and the LFP signal from the deep layers (600–800 *µ*m) was 0.35 (interquartile range: 0.32–0.38, n=4 animals) (Figure4C). The permutation test confirmed that this correlation was highly statistically significant (p ¡ 0.001). This result demonstrates that the flexible subdural array is capable of capturing neural activity that is representative of the underlying cortical processing.

## Discussion

In this paper, we presented a complete algorithm for the design, fabrication, and validation of a flexible subdural electrode array. The main advantage of our method is its accessibility and speed. By using a laser cutter for both mask creation and direct metal patterning on PDMS, we eliminate the need for expensive and labor-intensive cleanroom lithography for each design iteration. This rapid prototyping approach allows researchers to move from computer-aided design (CAD) to a functional device in significantly less time and at a lower cost than traditional microfabrication methods. [11]. The resolution of *∼*30 *µm* achieved by our laser ablation process is sufficient for fabricating macro- and micro-ECoG arrays suitable for a wide range of pre-clinical and intraoperative applications.

Our choice of materials—polydimethylsiloxane (PDMS) as a substrate and gold as conductors—has proven success in the field of flexible bioelectronics. PDMS has a Young’s modulus orders of magnitude lower than that of silicon, providing better mechanical conformity with soft nerve tissue and potentially reducing foreign body reactions [23]. Gold provides excellent conductivity and biocompatibility for chronic implantation [14]. The galvanic reinforcement step was crucial to ensure the mechanical strength of the traces, as evidenced by the minimal change in conductivity after repeated bending. (Figure2B). The slight reversible change in conductivity under deformation is likely due to microcracks in the metal film that heal upon relaxation, a phenomenon previously observed in metal films on polymer substrates.

The impedance characteristics of our electrodes (Figure3C) are suitable for recording field potentials (ECoG, LFP). While the impedance might be higher than that of bulk platinum electrodes, it is comparable to other polymer-based thin-film electrodes [11] and is sufficient for capturing high-quality signals, as demonstrated by our *in vivo* recordings.

The most critical result of this study is the functional validation *in vivo*. The statistically significant cross-correlation (median r=0.35, *p <* 0.001) between the surface ECoG signal from our array and the LFP from deep cortical layers (Figure4C) is a key finding. It demonstrates that our device is not merely recording noise or artifacts, but is capturing physiologically relevant signals that are coherent with the activity of the underlying neuronal populations. This level of correlation is expected, as the ECoG signal is generated by the summation of synaptic currents from a large neuronal pool, which is also reflected in the LFP. This proof-of-concept aligns with the goal of creating reliable neural interfaces for intraoperative monitoring, where real-time, high-quality ECoG is essential for mapping functional areas and guiding surgical resections.

The successful long-duration recordings (up to 12 hours) without signal degradation or adverse effects on the animals further support the robustness and safety of the device for acute applications. The ease of handling and repositioning on the cortical surface without inducing trauma underscores its potential for intraoperative use.

## Conclusion

We have successfully developed and validated a rapid and inexpensive process for fabricating flexible PDMS-based subdural electrode arrays. Using laser ablation to pattern the arrays, we can quickly make design changes without sacrificing precision. The resulting devices possess robust mechanical and electrical properties suitable for reliable signal recording. Their functionality was confirmed in vivo by recording statistically significant correlated neural activity using a commercial penetrating probe in the rat cerebral cortex. This technology holds great promise for intraoperative neuromonitoring and as a platform for developing custom neural interfaces in preclinical studies. Future work will focus on chronic implantation studies to assess long-term stability and biocompatibility, as well as increasing electrode density for mapping with higher spatial resolution.

The authors declare no conflict of interest. Suchkov D.S., Malyshev E.I., Vorobyov A.A., Mitin D.M, Mukhin I.S., Belousov V.V. are inventors on patent applications related to flexible subdural electrode array for intraoperative mapping of cortical activity. Mamleev A.R., Malyshev E.I., Sitdikova V.R., Silaeva V.M., Logashkin A.E., Kireev A.K. were employees at LIFT LLC at the time the study was performed.

## References

1. T. Aflalo et al. Decoding motor imagery from the posterior parietal cortex of a tetraplegic human. Science, 348(6237): 906–910, 2015.

2. A. Altuna et al. Su-8 based microprobes with integrated planar electrodes for enhanced neural depth recording. Biosensors and Bioelectronics, 37(1): 1–5, 2012.

3. Z. Bao, J. A. Rogers, and H. E. Katz. High-performance plastic transistors fabricated by printing techniques. Chemistry of Materials, 9(6): 1299–1301, 1997.

4. O. E. Beder and G. Eade. An investigation of tissue tolerance to titanium metal implants in dogs. Surgery, 39(3): 470–473, 1956.

5. S. J. Bensmaia and L. E. Miller. Restoring sensorimotor function through intracortical interfaces: Progress and looming challenges. Nature Reviews Neuroscience, 15(5): 313–325, 2014.

6. R. G. Bickford et al. Histologic changes in the cat’s brain after introduction of metallic and plastic coated wire used in electro-encephalography. Proceedings of the Staff Meetings of the Mayo Clinic, 32(1): 14–21, 1957.

7. S. H. Cho et al. Biocompatible su-8-based microprobes for recording neural spike signals from regenerated peripheral nerve fibers. IEEE Sensors Journal, 8(11): 1830–1836, 2008.

8. A. Czeschik et al. Fabrication of mea-based nanocavity sensor arrays for extracellular recording of action potentials. Physica Status Solidi A, 211(6): 1462–1466, 2014.

9. M. Dimaki et al. Fabrication and characterization of 3d micro- and nanoelectrodes for neuron recordings. Sensors, 10(11): 10339–10355, 2010.

10. A. M. Dymond et al. Brain tissue reaction to some chronically implanted metals. Journal of Neurosurgery, 33(5): 574–580, 1970.

11. M. T. Flavin et al. Rapid and low cost manufacturing of cuff electrodes. Frontiers in Neuroscience, 15:628778, 2021.

12. T.-M. Fu et al. Stable long-term chronic brain mapping at the single-neuron level. Nature Methods, 13(10): 875–882, 2016.

13. F. Garnier, R. Hajlaoui, A. Yassar, and P. Srivastava. All-polymer field-effect transistor realized by printing techniques. Science, 265(5179): 1684–1686, 1994.

14. L. A. Geddes and R. Roeder. Criteria for the selection of materials for implanted electrodes. Annals of Biomedical Engineering, 31(7): 879–890, 2003.

15. G. H. Gelinck et al. Flexible active-matrix displays and shift registers based on solution-processed organic transistors. Nature Materials, 3(2): 106–110, 2004.

16. P. M. George et al. Fabrication and biocompatibility of polypyrrole implants suitable for neural prosthetics. Biomaterials, 26(17): 3511–3519, 2005.

17. A. E. Hess et al. Development of a stimuli-responsive polymer nanocomposite toward biologically optimized, mems-based neural probes. Journal of Micromechanics and Microengineering, 21(5): 054009, 2011.

18. J. Hsu et al. Encapsulation of an integrated neural interface device with parylene c. IEEE Transactions on Biomedical Engineering, 56(1): 23–29, 2009.

19. G.-T. Hwang et al. In vivo silicon-based flexible radio frequency integrated circuits monolithically encapsulated with biocompatible liquid crystal polymers. ACS Nano, 7(5): 4545–4553, 2013.

20. A. Jackson and E. E. Fetz. Compact movable microwire array for long-term chronic unit recording in cerebral cortex of primates. Journal of Neurophysiology, 98(5): 3109–3118, 2007.

21. S. B. Jun et al. Low-density neuronal networks cultured using patterned poly-l-lysine on microelectrode arrays. Journal of Neuroscience Methods, 160(2): 317–326, 2007.

22. D.-H. Kim et al. Conducting polymers on hydrogel-coated neural electrode provide sensitive neural recordings in auditory cortex. Acta Biomaterialia, 6(1): 57–62, 2010.

23. D.-H. Kim et al. Dissolvable films of silk fibroin for ultrathin conformal bio-integrated electronics. Nature Materials, 9(6): 511–517, 2010.

24. D.-H. Kim et al. Epidermal electronics. Science, 333(6044): 838–843, 2011.

25. D.-H. Kim et al. Materials for multifunctional balloon catheters with capabilities in cardiac electrophysiological mapping and ablation therapy. Nature Materials, 10(4): 316–323, 2011.

26. D.-H. Kim et al. Electronic sensor and actuator webs for large-area complex geometry cardiac mapping and therapy. Proceedings of the National Academy of Sciences USA, 109(49): 19910–19915, 2012.

27. D.-H. Kim et al. Thin, flexible sensors and actuators as ‘instrumented’ surgical sutures for targeted wound monitoring and therapy. Small, 8(21): 3263–3268, 2012.

28. J.-H. Kim et al. Surface-modified microelectrode array with flake nanostructure for neural recording and stimulation. Nanotechnology, 21(8): 085303, 2010.

29. J.-M. Kim et al. Plateau-shaped flexible polymer microelectrode array for neural recording. Polymers, 9(12): 690, 2017.

30. R. Kim and Y. Nam. Gold nanograin microelectrodes for neuroelectronic interfaces. Biotechnology Journal, 8(2): 206–214, 2013.

31. K.-K. Lee et al. Polyimide-based intracortical neural implant with improved structural stiffness. Journal of Micromechanics and Microengineering, 14(1): 32, 2004.

32. S. M. Lee et al. Self-adhesive epidermal carbon nanotube electronics for tether-free long-term continuous recording of biosignals. Scientific Reports, 4:6074, 2014.

33. C. T. Lin et al. Development of wireless brain computer interface with embedded multitask scheduling and its application on real-time driver’s drowsiness detection and warning. IEEE Transactions on Biomedical Engineering, 55(5): 1582–1591, 2008.

34. C. T. Lin et al. Noninvasive neural prostheses using mobile and wireless eeg. Proceedings of the IEEE, 96(7): 1167–1183, 2008.

35. K. A. Ludwig et al. Chronic neural recordings using silicon microelectrode arrays electrochemically deposited with a poly(3,4-ethylenedioxythiophene) (pedot) film. Journal of Neural Engineering, 3(1): 59–70, 2006.

36. K. Mathieson et al. Large-area microelectrode arrays for recording of neural signals. IEEE Transactions on Nuclear Science, 51(5): 2027–2031, 2004.

37. N. G. Page and M. A. Gresty. Motorist’s vestibular disorientation syndrome. Journal of Neurology, Neurosurgery & Psychiatry, 48(8): 729–735, 1985.

38. D.-W. Park et al. Fabrication and utility of a transparent graphene neural electrode array for electrophysiology, in vivo imaging, and optogenetics. Nature Protocols, 11(11): 2201–2222, 2016.

39. S. Park et al. Nanoporous pt microelectrode for neural stimulation and recording: In vitro characterization. The Journal of Physical Chemistry C, 114(19): 8721–8726, 2010.

40. J. A. Perge et al. Intra-day signal instabilities affect decoding performance in an intracortical neural interface system. Journal of Neural Engineering, 10(3): 036004, 2013.

41. V. S. Polikov, P. A. Tresco, and W. M. Reichert. Response of brain tissue to chronically implanted neural electrodes. Journal of Neuroscience Methods, 148(1): 1–18, 2005.

42. R. H. Reuss et al. Macroelectronics: Perspectives on technology and applications. Proceedings of the IEEE, 93(7): 1239–1256, 2005.

43. F. R. Robinson and M. T. Johnson. Histopathological studies of tissue reactions to various metals implanted in cat brains. ASD Technical Report, 61:13, 1961.

44. J. A. Rogers et al. Paper-like electronic displays: Large-area rubber-stamped plastic sheets of electronics and microencapsulated electrophoretic inks. Proceedings of the National Academy of Sciences USA, 98(9): 4835–4840, 2001.

45. J. A. Rogers, T. Someya, and Y. Huang. Materials and mechanics for stretchable electronics. Science, 327(5973): 1603–1607, 2010.

46. R. Saha et al. Highly doped polycrystalline silicon microelectrodes reduce noise in neuronal recordings in vivo. IEEE Transactions on Neural Systems and Rehabilitation Engineering, 18(5): 489–497, 2010.

47. E. Seker et al. The fabrication of low-impedance nanoporous gold multiple-electrode arrays for neural electrophysiology studies. Nanotechnology, 21(12): 125504, 2010.

48. J. H. Shin et al. Carbon-nanotube-modified electrodes for highly efficient acute neural recording. Advanced Healthcare Materials, 3(2): 245–252, 2014.

49. D. M. Simon et al. Design and demonstration of an intracortical probe technology with tunable modulus. Journal of Biomedical Materials Research Part A, 105(1): 159–168, 2017.

50. D. Suchkov et al. Weegit – software for visualization and annotation of the electrophysiological activity registration data. Zhurnal Vysshei Nervnoi Deyatelnosti Imeni I.P. Pavlova, 73: 857–872, 2023.

51. Y. Takayama et al. Network-wide integration of stem cell-derived neurons and mouse cortical neurons using microfabricated co-culture devices. BioSystems, 107(1): 1–8, 2012.

52. J. Viventi et al. A conformal, bio-interfaced class of silicon electronics for mapping cardiac electrophysiology. Science Translational Medicine, 2(24):24ra22, 2010.

53. J. Viventi et al. Flexible, foldable, actively multiplexed, high-density electrode array for mapping brain activity in vivo. Nature Neuroscience, 14(12): 1599–1605, 2011.

54. C. Xie et al. Intracellular recording of action potentials by nanopillar electroporation. Nature Nanotechnology, 7(3): 185–190, 2012.

55. B. Xu et al. An epidermal stimulation and sensing platform for sensorimotor prosthetic control, management of lower back exertion and electrical muscle activation. Advanced Materials, 28(22): 4462–4471, 2016.

56. A. Zátonyi et al. Functional brain mapping using optical imaging of intrinsic signals and simultaneous high-resolution cortical electrophysiology with a flexible, transparent microelectrode array. Sensors and Actuators B: Chemical, 273: 519–526, 2018.

